## Supplementary Figures for "A molecular proximity sensor based on an engineered, dual-component guide RNA"

**Supplementary materials for: A molecular proximity sensor based on an engineered, dual-component guide RNA**

List of Supplementary Figures:

1. Testing the sgRNA:petRNA splitting strategy.
2. Testing the self-splicing ribozyme strategy.
3. Characterizing linkers for crRNA-MS2 and BoxB-petracrRNA.

List of Supplementary Data:

1. Nucleic acid sequences used in the study
2. Numbers underlying editing efficiency data shown in the figure

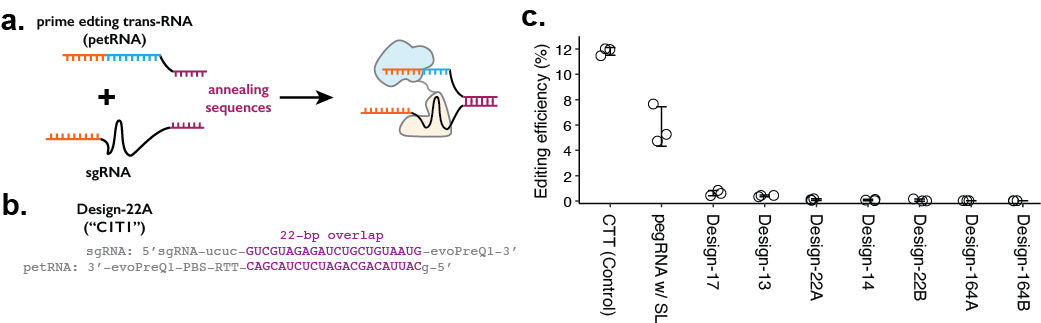

#### Supplementary Figure 1. Testing the sgRNA:petRNA splitting strategy.

**a.** Schematic of splitting functional pegRNA into sgRNA with extended sequence on 3’-end and petRNA with extended sequence on 5’-end for controlling dimerization. Annealing of two dimerization sequences would result in an active pegRNA. **b.** Example design of dimerization sequence. Each design is named after the length of the reverse-complementary sequence (22-bp shown here). In both sgRNA and petRNA, evoPreQ1 RNA pseudoknot domains are added to reduce the degradation of the non-Cas9-bound RNA portion. **c.** The editing efficiency was measured for seven different sgRNA:petRNA pairs along with prime editing control (pegRNA targeting HEK3 locus for CTT insertion at +0 position from the nick) and a single pegRNA construct with 22-bp RNA stem-loop between the sgRNA and RTT portion (pegRNA w/ SL) to test whether having extra RNA-duplex affects the prime editing efficiency. The center and error bars are mean and standard deviations, respectively, from n = 3 transfection replicates.

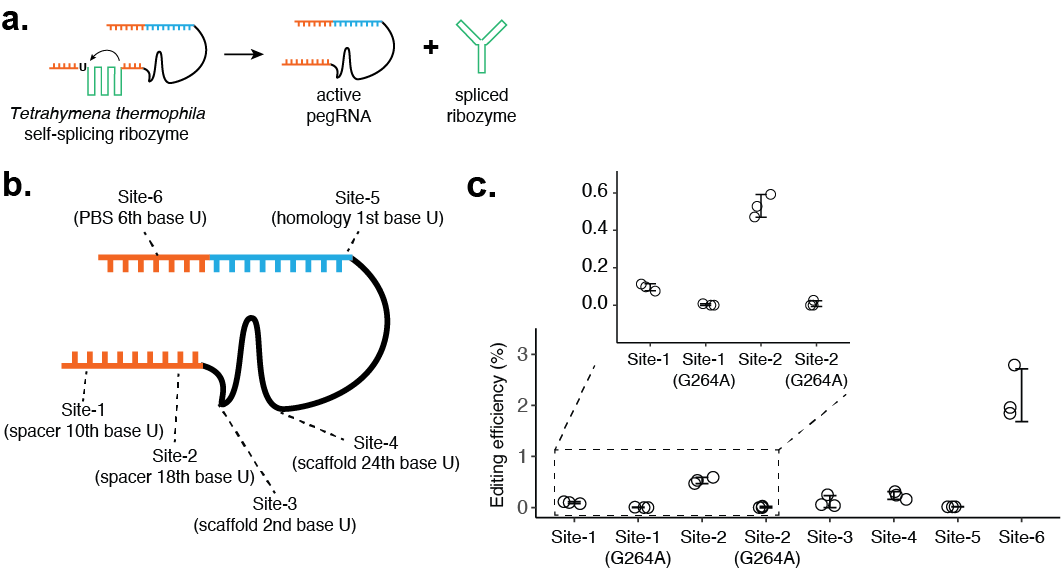

#### Supplementary Figure 2. Testing the self-splicing ribozyme strategy.

### **a.** Schematic of self-splicing ribozyme strategy. Insertion of the ribozyme into functional pegRNA would lead to self-splicing after RNA transcription, which would result in an active pegRNA and spliced ribozyme sequence. If this worked, the ribozyme could be further split into two functional parts, where extra dimerization domains can be added to control ribozyme function. **b.** Six positions within the pegRNA (targeting HEK3 locus for CTT insertion) were tested for inserting the self-splicing ribozyme sequence. Ribozyme needs to start with a uridine base for splicing, which remains after the ribozyme is spliced out. **c.** Editing efficiencies were measured for 6 ribozyme insertion sites within pegRNA. For Site-1 and Site-2, two additional designs were constructed with inactive ribozyme (G264A mutation). The center and error bars are mean and standard deviations, respectively, from n = 3 transfection replicates.

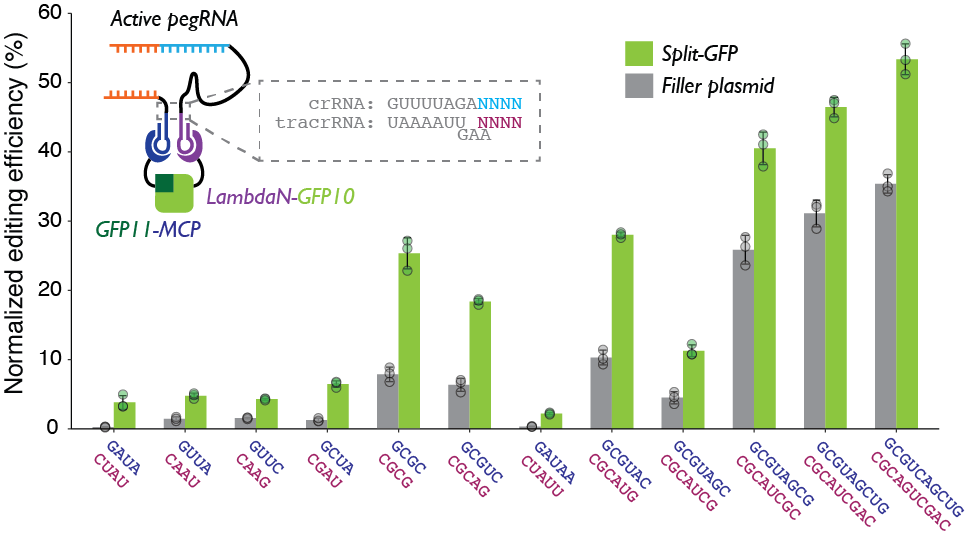

#### Supplementary Figure 3. Characterizing linkers for crRNA-MS2 and BoxB-petracrRNA.

The upper four base pairs of the Cas9-binding region of repeat:anti-repeat duplex were altered to generate 12 pairs of crRNA-MS2/BoxB-petracrRNA designs. The editing efficiency is normalized with the eCTT positive control included in each experiment (targeting HEK3 locus with CTT insertion at position +0 using standard epegRNA), to control for variable transfection efficiencies. Two normalized editing efficiencies were measured for each pair of RNAs: one with tagged split-GFP to promote dual-RNA-guide formation, and one with standard, untagged GFP to measure the background editing level non-specific to protein-protein proximity. The center and error bars are mean and standard deviations, respectively, from n = 3 transfection replicates.
